# Towards resilient beef cattle production systems: Impact of truck contamination and information sharing on foot-and-mouth disease spreading

**DOI:** 10.1101/2020.04.17.046789

**Authors:** Qihui Yang, Don M. Gruenbacher, Jessica L. Heier Stamm, David E. Amrine, Gary L. Brase, Scott A. DeLoach, Caterina M. Scoglio

## Abstract

As cattle movement data in the United States are scarce due to the absence of mandatory traceability programs, previous epidemic models for U.S. cattle production systems heavily rely on contact rates estimated based on expert opinions and survey data. These models are often based on static networks and ignore the sequence of movement, possibly overestimating the epidemic sizes. In this research, we adapt and employ an agent-based model that simulates beef cattle production and transportation in southwest Kansas to analyze the between-premises transmission of a highly contagious disease, the foot-and-mouth disease. First, we assess the impact of truck contamination on the disease transmission with the truck agent following an independent clean-infected-clean cycle. Second, we add an information-sharing functionality such that producers/packers can trace back and forward their trade records to inform their trade partners during outbreaks. Scenario analysis results show that including indirect contact routes between premises via truck movements can significantly increase the amplitude of disease spread, compared with equivalent scenarios that only consider animal movement. Mitigation strategies informed by information sharing can dramatically improve the system resilience against epidemics, highlighting the benefit of promoting information sharing in the cattle industry. In addition, we identify salient characteristics that must be considered when designing an information-sharing strategy, including the number of days to trace back and forward in the trade records and the role of different cattle supply chain stakeholders. Sensitivity analysis results show that epidemic sizes are sensitive to variations in parameters of fomite survival time and indirect contact transmission probability and future studies can focus on a more accurate estimation of these parameters.

## 1. Introduction

Foot-and-mouth disease (FMD) is a highly contagious infectious disease that could threaten cloven-hoofed animals worldwide [1]. The FMD outbreak in 2001 in the United Kingdom resulted in severe economic losses totaling more than £2.8 billion and required the slaughter of approximately 6.5 million animals [2]. Recently, several major outbreaks have occurred in previously FMD-free countries, including South Korea, Japan, and Uganda [3–5]. Although the United States has been free of FMD since 1929, the disease is still prevalent in approximately two-thirds of the world [6]. Concerns about the reintroduction of FMD into the United States have escalated due to increases in international travel and trade [7]. The beef cattle industry is of great importance to the U.S. economy, and an FMD outbreak would produce devastating economic losses as estimated by simulation-based studies. For example, Pendell et al. [8] reported that an FMD virus released from the National Bio and Agro-Defense Facility in Kansas could cause $16 billion– $140 billion in damages. Schroeder et al. [9] estimated that a hypothetical FMD outbreak in the midwestern United States could result in $56 billion–$188 billion of producer and consumer losses.

The FMD virus can be transmitted between farms through animal movement (direct contact) and via fomites such as contaminated equipment and vehicles (indirect contact) [10–12]. While disease transmission through the movement of infected cattle has been extensively studied in previous works [13–15], indirect contact through fomites has recently gained attention, especially after the 2001 FMD outbreak in the United Kingdom, in which new cases occurred for several months after early implementation of an animal movement ban [16]. The spread of FMD stopped only after strict biosecurity measures targeting the movement of contaminated equipment and personnel were implemented [17]. This study focused on indirect contact via livestock transporters, one of the most at-risk operator categories [10,18].

Only a few studies have considered the truck movement to be indirect contact routes for the spread of disease among farms. Thakur et al. [19] developed a farm-level model to simulate the spread of porcine reproductive and respiratory syndrome virus through animal movement and truck sharing. Their results of using static links between farms highlighted the significant role indirect contact played in spreading the disease. Wiltshire [20] developed a model to heuristically generate a dynamic hog production system while accounting for truck contamination, demonstrating that producer specialization can increase system vulnerability to disease outbreaks. Bernini et al. [2] developed a two-layer temporal network to model disease transmission through cattle exchanges and transportation and compared epidemic size under full or partial knowledge of daily truck itineraries. Their work concluded that an accurate description of indirect contact is essential for precise prediction of epidemic spreading dynamics. However, no previous studies have included trucks as independent epidemiological units, as is emphasized in this research.

Simulation models have been widely used to mimic FMD transmission among farms and analyze various control strategies. Bate et al. [7] simulated FMD transmission in a three-county area in California, demonstrating the effectiveness of preemptive culling of highest-risk herds and ring vaccination. This model was widely used and later adapted to analyze FMD mitigation strategies in other countries, including Sweden [21] and Denmark [11]. Other stochastic, state-transition simulation models included the North American Animal Disease Spread Model [22], AusSpread [23], and InterSpreadPlus [24], which involve multiple animal species and pathways for disease spreading. However, these herd-level models rely on accurate estimates of the contact frequency and distance distribution between livestock operations and do not consider actual contact sequences that occur between farms [2]. In addition, AusSpread does not explicitly model disease spread within a farm. Although the Australian Animal Disease model [25] predicts the fraction of animals in each state of each herd, the number of animals is simplified to be constant over time for each herd agent.

Suggested animal movement traceability systems in the United States have faced opposition from beef producer organizations due to privacy issues, resulting in contact rates used for most FMD simulation models being based on expert opinions and questionnaires. Though the inspection of slaughterhouses is the most effective surveillance component for early epidemic detection [26], most FMD simulation models are built without including slaughterhouses. In addition, they are often based on static networks and ignore the system’s time-varying structure, which is crucial for simulating the dynamics of highly contagious diseases [27]. Recent work has focused increasingly on temporal network measures, showing that possible outbreak sizes may be overestimated in a static view of the network [28]. Sterchi et al. [29] concluded that information about transport sequences could change the contact network topology and that consideration of truck sharing and contamination could increase network connectivity and individual connectedness of farms. Liu et al. [30] built a spatially explicit cattle-level agent-based model for two counties in Kansas, in which producers made decisions on cattle trade based on cattle weights and market conditions. Their work emphasized the influence of trading dynamics on the disease transmission through cattle movement.

Traceability programs, however, have become common in the global beef market, and lack of such programs may decrease export markets for the U.S. beef industry [31]. For example, if 25% of beef products became unacceptable in international trade, then the U.S. economy would experience an estimated $6.65 billion loss [32]. An improved information infrastructure with traceability systems would yield many benefits, including targeted and timely product recalls after a foodborne illness outbreak and increased brand value for products due to quality assurance. Several pilot projects have recently developed and tested purpose-built cattle disease traceability infrastructures, such as BeefChain and CattleTrace. In these projects, cattle movement information is uploaded to a secure third-party database through radio frequency identification tags or similar devices. Since the acceptance of such traceability programs depends on the trust among stakeholders, these pilot programs have focused on persuading operators to participate. New technologies such as Blockchain, which has been increasingly studied [33–36], are expected to build trust among food partners, promote livestock traceability, and enhance food safety.

In response to potential FMD outbreaks, governmental agencies in the United States have designed control measures and preparedness plans. According to the U.S. Department of Agriculture, the depopulation of clinically affected and in-contact susceptible animals (stamping-out) is typically applied with other emergency vaccination strategies depending on the circumstances and epidemic sizes of FMD outbreaks [6]. In Kansas, for example, the Kansas Animal Health Commissioner would issue a state-wide stop movement order to all animal and related product movement if an FMD case occurred in North America. This movement restriction would remain in effect until the situation was deemed safe for the Kanas livestock industry, and producers would have to provide documents such as normal health status for animals on the production site for the previous 14 days to request a movement permit from the Kansas Department of Agriculture [37].

This work aims to *examine the potential impact of truck contamination and information sharing for FMD virus transmission in the beef cattle production system in southwest Kansas* (SW KS), *United States*. We simulate a hypothetical FMD transmission through cattle movement and truck movement with major expansions relative to the model in [38], including state transition models embedded in cattle agent, truck agent, producer agent, and packer agent. In order to enable information-sharing functionality, each producer/packer agent stores time-stamped trade information and can inform their trade partners during an outbreak. Scenario analysis is conducted to evaluate the effect of various control strategies on the spread of the epidemic, and sensitivity analysis is implemented to examine the outcomes affected by different input parameters.

This study does not mean to predict the spread of FMD in a real-world production system, but it facilitates a realistic epidemiological model to highlight the impact of indirect contact through truck movement and the potential benefits of information sharing to the system regarding disease transmission. The findings are expected to benefit existing disaster preparedness and promote the development of new mitigation strategies informed by information sharing for rapid detection and containment.

## 2. Materials and methods

In this section, we introduce the major expansions to a previously developed agent-based model and briefly describe the experimental design.

### 2.1 Epidemiological model

In this work, an agent-based, stochastic, cattle-level simulation model is developed in AnyLogic software. Due to the model’s large complexity, we have described only the added features to the base model in [38], which generates dynamic cattle movement and truck movement networks based on regular business operating principles and assumed conditions. Agent functionalities are altered to enable FMD transmission through both direct and indirect contact routes and information-sharing functionality.

#### 2.1.1 Study area

The study area is SW KS, consisting of 24 counties. As shown in Fig 1, cattle producers include cow-calf ranches, stockers, and feedlots. In ranches, cows produce a new generation of calves that are fed until around 450 pounds and that will be sold directly or via auction markets to stockers. At approximately 650 pounds, cattle raised in stocker operations will be sold directly or via auction markets to feedlots. Once heifers and steers in feedlots achieve 1250 pounds and 1350 pounds respectively, they will be moved to the final meat-processing facilities, the packers. When producers from inside SW KS request cattle from the outside, cattle coming from outside SW KS are generated at entry point agents, which are located on the major highways into and out of SW KS. The study population and parameters used for simulation are presented in Table 1.

**Table 1.**
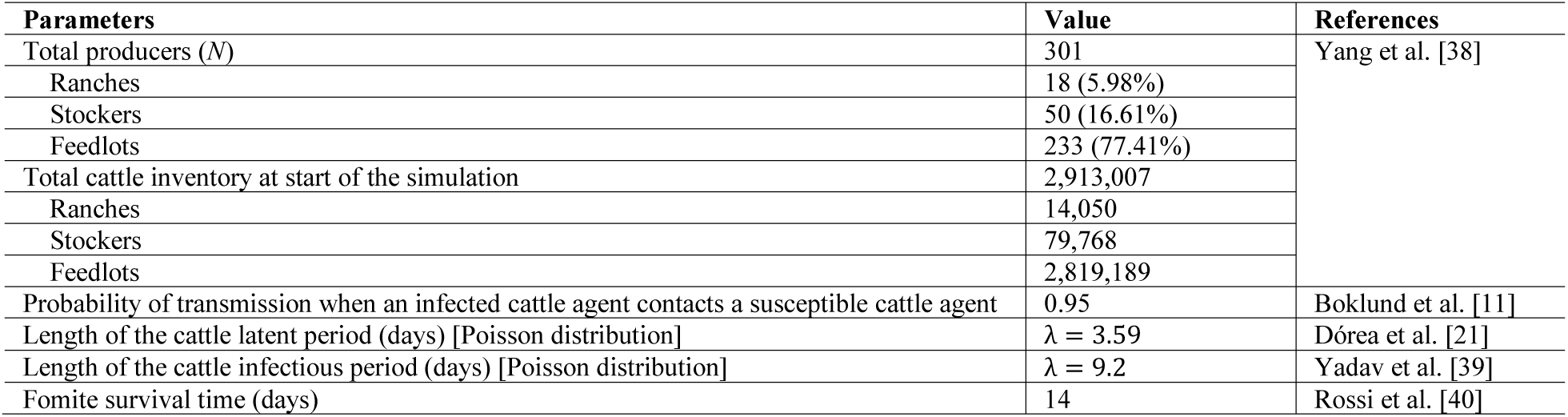
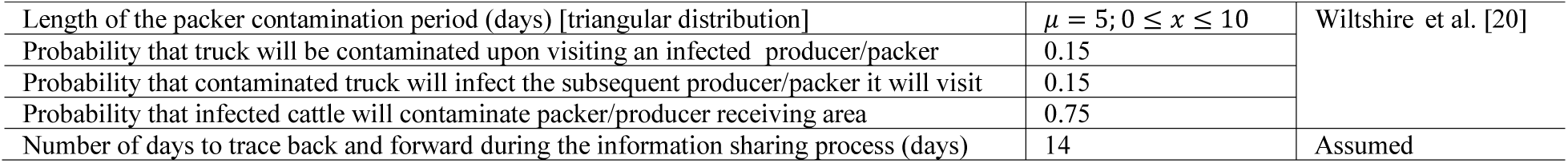
Study population and parameters used for simulation of FMD spread.

**Fig 1.**
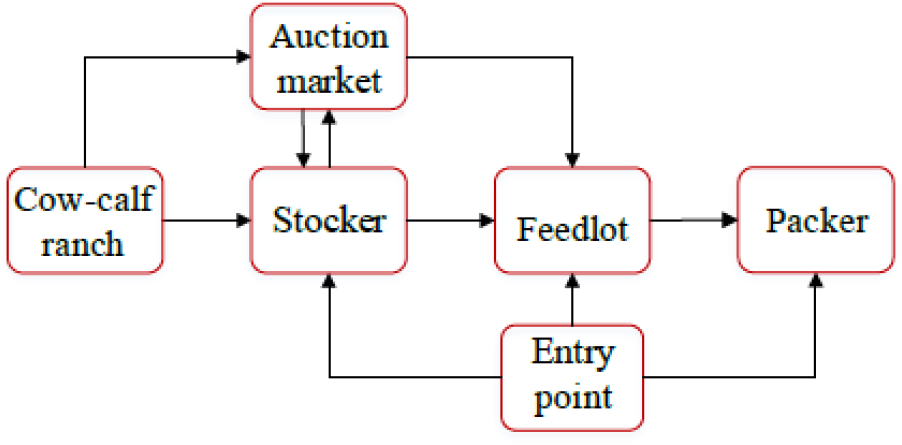
Cattle flow through the system.

#### 2.1.2 Epidemic initialization

All the cattle in the system are in the susceptible state before the epidemic is initialized. For each simulation run, one randomly selected animal from outside SW KS will enter the infectious state on the 9^th^ day midnight, and will be brought to a stocker inside the region on the 10^th^ day. If there are no cattle at the border at this time, the simulation will stop and start the next simulation run. In different simulation runs, the recipient stocker of the first infectious animal will be different due to the stochasticity of the model.

#### 2.1.3 Spread of the disease

In this section, we describe major functions added to the base model to enable FMD transmission using Fig. 2.

**Fig 2.**
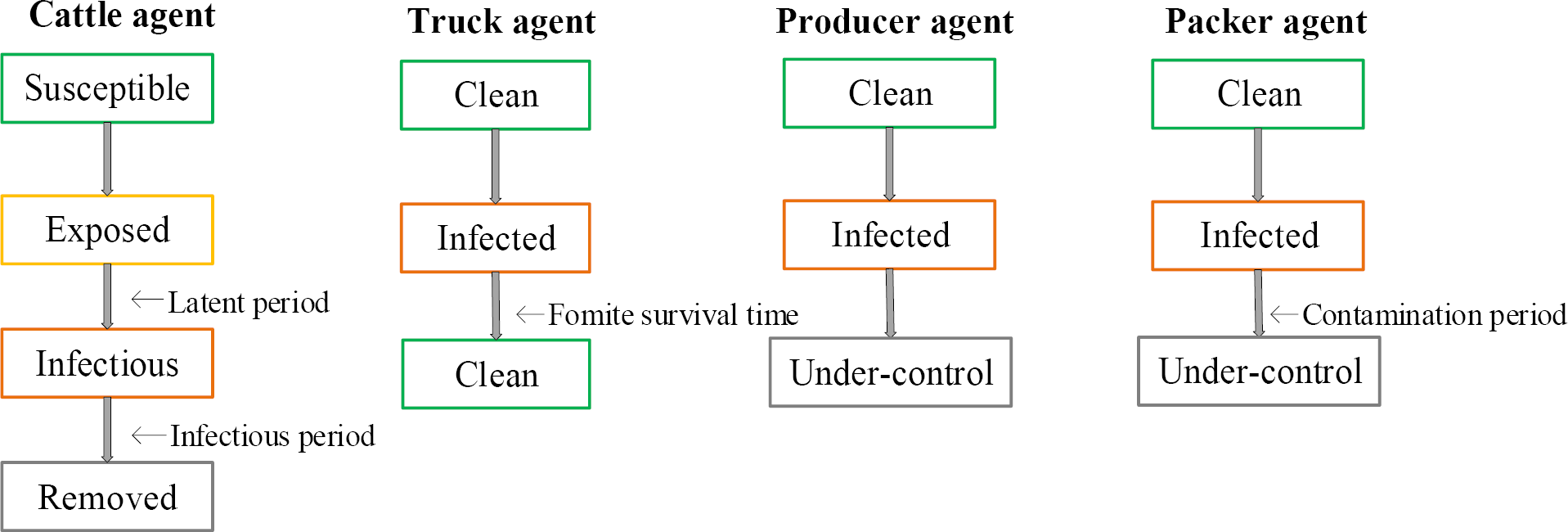
Transition diagram for agents added with epidemiological components.

##### (1) Cattle agent

Each cattle agent is associated with a Susceptible-Exposed-Infectious-Removed compartment model. Compared to the cattle in the infectious state, cattle agents in the exposed state have been infected by the FMD disease, but are not contagious yet. Animals in the removed state have been infected before, and become dead or immune, or are culled as part of control strategies. More specifically, in contact with an animal in the infectious state, a susceptible cattle agent has a 95% chance to become infected and transition to the exposed state. After a latent period, the cattle agent will transition to the infectious state during which it contacts and infects other cattle of the same premises. Then, the cattle agent enters the removed state after an infectious period.

Considering computational efficiency, we use a scaling factor of 10 to change all the parameters related to the number of cattle, e.g., truck capacity and cattle capacity of each producer, such that one cattle agent represents ten cattle. We assume that each cattle agent contacts, on average, 20 cattle agents in a day. 20 cattle agents correspond to 200 cattle due to the scaling factor, indicating the approximate number of cattle in a group. Accordingly, during the infectious state, each cattle agent can infect on average 20*0.95 cattle agents per day, where 0.95 is the probability of transmission when infected cattle contact susceptible cattle.

##### (2) Producer agent

Once there are infectious cattle in the producer agent, the producer will transition from the clean state to the infected state and the *c_infected* variable becomes true. After a cattle infectious period, the first infectious cattle agent will become removed, and if control strategies have been implemented on the producer premises level, the producer agent will enter the under-control state. In the under-control state, various mitigation strategies are implemented including movement bans and information-sharing functionality, and more details will be described in the scenario analysis section. When all infected cattle of an infected producer enter the removed state, the *c_infected* variable is set to be false. On the other hand, when a producer becomes infected by fomite, the variable *f_infected* becomes true and will last for 14 days (fomite survival time in Table 1).

##### (3) Packer agent

Once a cattle agent in the infectious state arrives at the packer, the packer agent will transition to the infected state (c*_infected*=true), and after a contamination period *T*, the packer goes to the under-control state, in which the packer stops requesting and transporting cattle from other producers to its location. To be more realistic, we assume that more infectious cattle arriving at the packer will speed up FMD detection. For example, if there are infectious cattle coming in on day *d*1 and day *d*2, the model will generate two numbers, *ct* and *ct*’, respectively, according to the distribution with a mean of 5 days in Table 1. The smaller value between *ct*’ and *ct* will be assigned as the contamination period *T*, as shown in Fig 2. On the other hand, once the packer is infected by fomite, its cattle receiving area will remain contaminated (*f_infected* = true) for 14 days based on the fomite survival time.

##### (4) Truck agent

Following a clean-infected-clean cycle, trucks move to the origin premises to pick up cattle and then move the animals to the destination premises to unload. With direct contact infection probability set as 1.0 [19,40], when at least one infectious cattle agent is moved and received at the destination premises, the destination becomes infected with *c_infected* set as true. For indirect contact, trucks may become contaminated by visiting fomite-infected premises, and then remain contaminated for a fomite survival period, during which time the infection may spread to other premises.

More specifically, when the truck arrives at the origin premises, if there are infectious cattle loaded to the truck at the origin premises, then there is a 75% probability that the receiving area of the origin producer will become infected via fomite (*f_infected* = true). If the origin producer is fomite-infected and the truck is not contaminated, the truck may become contaminated with a 15% probability; however, if the truck is contaminated and the origin producer is not fomite-infected (*f_infected* = false), the truck will cause the producer to become fomite-infected with a 15% probability.

When the truck arrives at the destination premises, if there are infectious cattle unloaded, there is a 75% probability that the destination premises will become fomite-infected. Meanwhile, if the destination is fomite-infected and the truck is clean, there is a 15% probability that the truck may become contaminated. However, if the truck is infected and the destination is not fomite-infected, the truck may cause the destination to become fomite-infected. Note that at the time the origin or destination producer becomes fomite-infected, we will randomly select one cattle agent from the premises to become infected.

### 2.2 Scenario analysis

In this section, ten different scenarios are constructed based on three attributes: (1) the two FMD virus transmission routes, namely by direct contact only or by both direct and indirect contact; (2) the implementation of movement bans; and (3) the involvement of information infrastructure. The combination of these factors is described in Table 2, and all the parameters follow the values described in Table 1 in all baseline scenarios. Three hundred iterations of each scenario are run to generate a distribution of the outcomes. A 200-day simulation duration is selected such that all producers already reach the steady-state in the end regarding disease transmission, i.e., no new infections occur.

**Table 2.** Description of the scenarios

| Scenario number | Scenario name | Spread by direct contact | Spread by indirect contact | Movement ban on infected producers | Information infrastructure enabled on producers | Information infrastructure enabled on packers | Movement ban in the whole region |
| --- | --- | --- | --- | --- | --- | --- | --- |
| 1 | DI | Yes | No | No | No | No | No |
| 2 | DI_B | Yes | No | Yes | No | No | No |
| 3 | DI_B_P | Yes | No | Yes | Yes | No | No |
| 4 | DI_B_PP | Yes | No | Yes | Yes | Yes | No |
| 5 | DI_RB | Yes | No | No | No | No | Yes |
| 6 | D&IN | Yes | Yes | No | No | No | No |
| 7 | D&IN_B | Yes | Yes | Yes | No | No | No |
| 8 | D&IN_B_P | Yes | Yes | Yes | Yes | Yes | No |
| 9 | D&IN_B_PP | Yes | Yes | Yes | Yes | Yes | No |
| 10 | D&IN_RB | Yes | Yes | Yes | No | No | Yes |

DI: direct contact only, D&IN: direct and indirect contact, B: movement ban on infected producers, P: information infrastructure on producers, PP: information infrastructure on both producers and packers. RB: region-wide movement ban.

In scenarios 1 and 6, there is no control strategy implemented on the producer premises level, and thereby the producers will not transition to the under-control state. In all the other scenarios, when an infectious cattle agent occurs in the producer, the producer becomes infected (*c_infected* = true). After an infectious period, the first infectious cattle agent becomes removed, and the infected producer is under control. In addition, all cattle in these infected producers under control will transition to the removed state.

More specifically, for scenarios considering movement restrictions, we assume that any movement ban was 100% effective, meaning that all movements to and from the producer would stop, once the movement ban is enacted. (i) For scenarios with a movement ban on infected producers, or called producer isolation (scenarios 2 and 7), a movement ban will be introduced to the infected producer once the producer is under control. (ii) For scenarios with a region-wide movement ban (scenarios 5 and 10), the movement ban will be enacted for all the producers and packers inside the region as long as there is one infected producer under control.

To enable information-sharing functionality, each producer/packer agent stores time-stamped trade information with its trade partners. During outbreaks, the producer/packer agent can trace back and forward its trade record in the past 14 days to inform their trade partners about its infection by disease. This time-interval of 14 days will be referred as the number of days to trace back and forward in the sensitivity analysis section. (i) If information-sharing functionality is enabled only on producers and not on packers (scenarios 3 and 8), the producer in the under-control state will notify those producers with which it has traded in the past 14 days (both its suppliers and customers). Once those producers receive notification from the information sender, they will go to the under-control state and notify their trading partners as well. (ii) For scenarios that information-sharing functionality is enabled on both producers and packers (scenarios 4 and 9), the producer will notify both producers and packers with which it has traded in the past 14 days. In addition, once an infected packer enters the under-control state, it will immediately notify its trade partner producers. On the other hand, if a producer in the under-control state notifies a packer not in the under-control state that it has the potential to be infected, the packer will also go to the under-control state. Since those producers may have cattle in exposed state at the time when they are informed by trade partners and transition to the under-control state, all their cattle will be culled (enter the removed state) once infectious cattle occur.

### 2.3 Sensitivity analysis

To evaluate how changes in parameters can impact the simulation results, we perform sensitivity analyses for different scenarios by creating variations in a single input parameter while keeping other simulation settings unchanged. First, three variations to the baseline scenarios D&IN_B_PP (information infrastructure enabled on producers and packers) and D&IN_B_P (information infrastructure enabled on only producers) are created with different values of the number of days to trace back and forward during the information-sharing process. The number of days to trace back and forward is selected for sensitivity analysis, as it is assumed as 14 days based on expert opinion and it is likely to influence the epidemic control effectiveness during the information-sharing process. Second, parameters related to indirect contact routes: (a) indirect contact transmission probability, and (b) fomite survival time are altered, and their impact on the number of infected producers are evaluated under scenario D&IN_B. In this work, we set indirect contact transmission probability as 0.15 based on [20], but Bate et al. [7] assume distributions for the high-risk and low-risk indirect contact probability as BetaPert(0.1,0.5,0.9) and BetaPert(0.05,0.175,0.35), respectively. Accordingly, we modify the baseline indirect contact transmission probability to more values ranging from 0.05 to 0.5 and compare the number of infected producers with the baseline output. In addition, we alter the baseline fomite survival time 14 days as previously assumed in [40] ranging from 1 day to 21 days, and compare the number of infected producers with the baseline scenario. Third, since there is variability in cattle infectious period in references, we evaluate the impact of varying the baseline cattle infectious period 9.2 days for scenarios DI_B (direct contact only) and D&IN_B (direct and indirect contact). For all variations, 300 iterations of each scenario are simulated for 200 days.

## 3. Results and discussion

### 3.1 Scenario analysis

Fig. 3 shows the distributions of numbers of infected producers and cattle agents removed for the ten scenarios. Overall, the epidemic size is smaller for scenarios that model only direct contact (scenarios 1–5) when compared to equivalent scenarios that incorporate both direct and indirect contact (scenarios 6–10). For example, the median number of infected producers changed from 14 in scenario 1 (direct contact only) to 90 in scenario 6 (direct and indirect contact). Findings indicate that indirect contact through truck contamination has a considerable impact on the FMD spread among producers. Since contaminated trucks can travel to multiple places, truck contamination can aggravate the disease spread between those who do not have direct contact (animal movement in between), thereby enlarging the scale of epidemic spreading.

**Fig 3.**
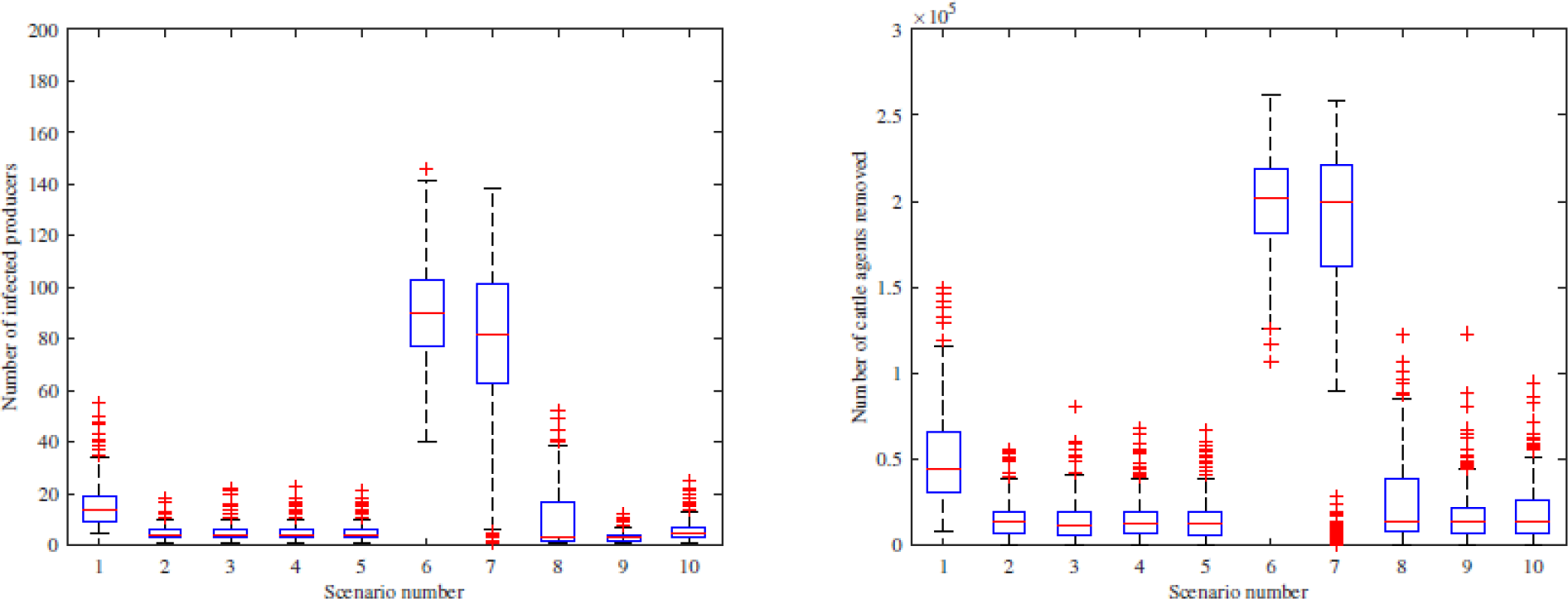
Distributions of numbers of infected producers (left) and cattle agents removed (right).

More specifically, for scenarios 1–5, scenarios 2–5 result in a smaller and similar epidemic size compared to scenario 1 (no premises-level control strategies implemented). For scenarios 6–10, scenario 6 (no premises-level control strategies implemented) results in the largest epidemic size, followed by scenario 7 (producer isolation), and then scenario 8 (information infrastructure enabled on producers), while scenarios 9 (information infrastructure enabled on producers and packers) and 10 (regional movement ban) result in the smallest and similar epidemic sizes. There is a slight difference in terms of the median size between scenarios 6 and 7, indicating that merely implementing movement bans on infected producers cannot effectively contain the epidemic, when indirect contact is considered. Information sharing can significantly reduce the epidemic size, compared to scenario 6. Particularly, scenario 8 has a similar median number of infected producers with scenarios 9 and 10 but has a larger interquartile range with 15 as the third quartile in Fig. 3. This is because information-sharing functionality is not enabled on packers in scenario 8, and these infected packers can spread the infection by their contaminated trucks to other producers in some simulation runs.

As additional information for the epidemic dynamics, Figs. 4–5 show the numbers of cattle agents in each compartment and newly infected producers over time for scenarios 2, 7 and 8–9. In Fig. 4, both the epidemic size and epidemic duration is larger in scenario 7 compared with scenario 2, indicating that truck contamination can aggravate the spread of epidemics. Comparing Fig. 5 with Fig. 4, it is shown that the addition of information-sharing functionality reduces the number of cattle agents removed during the outbreak. Particularly, there is a larger variance in terms of new infected producers in scenario 8 compared to scenario 9, indicating that the system is more robust when information-sharing functionality is enabled in both producers and packers.

**Fig 4.**
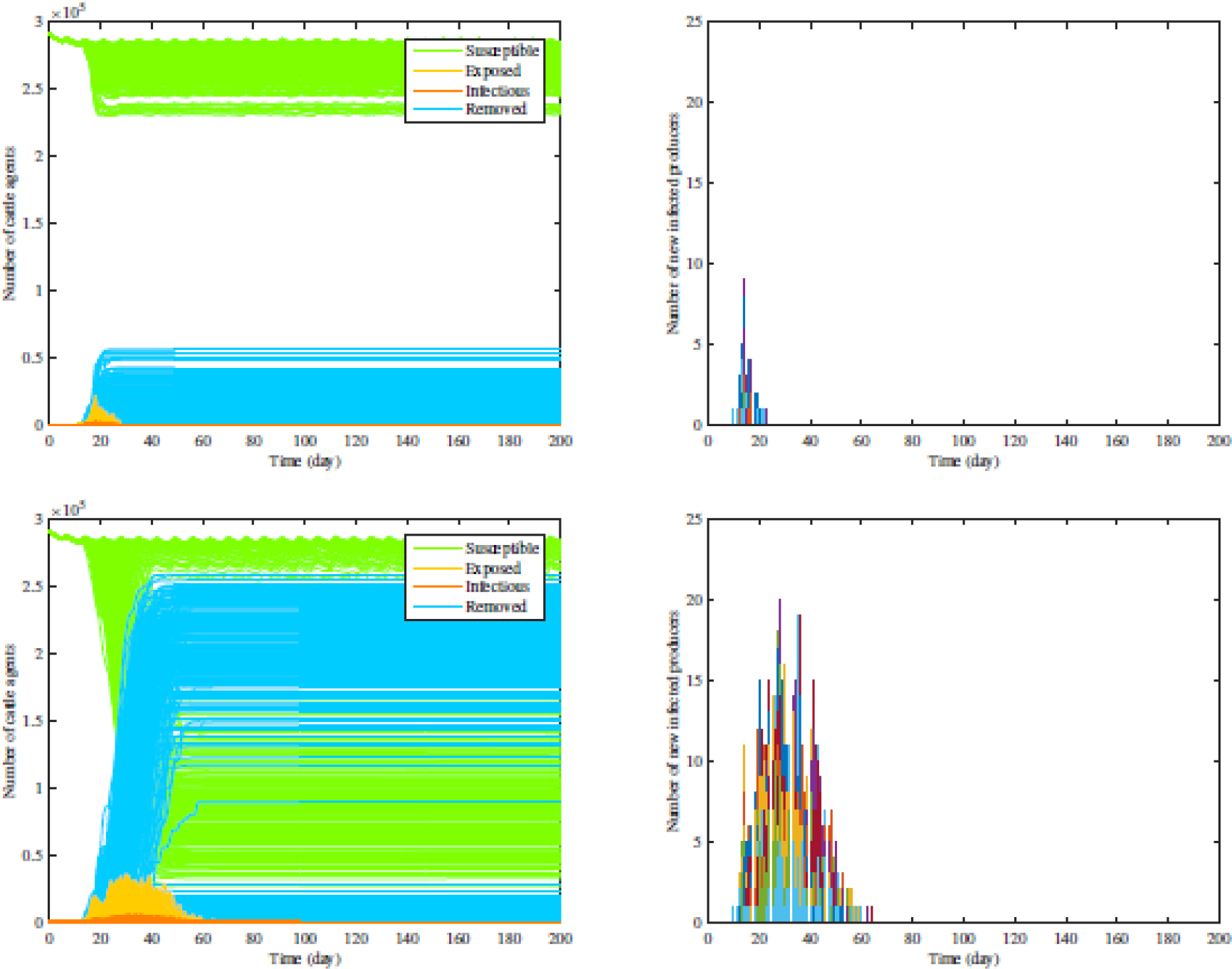
Comparison between scenarios 2 (top) and 7 (bottom).

**Fig 5.**
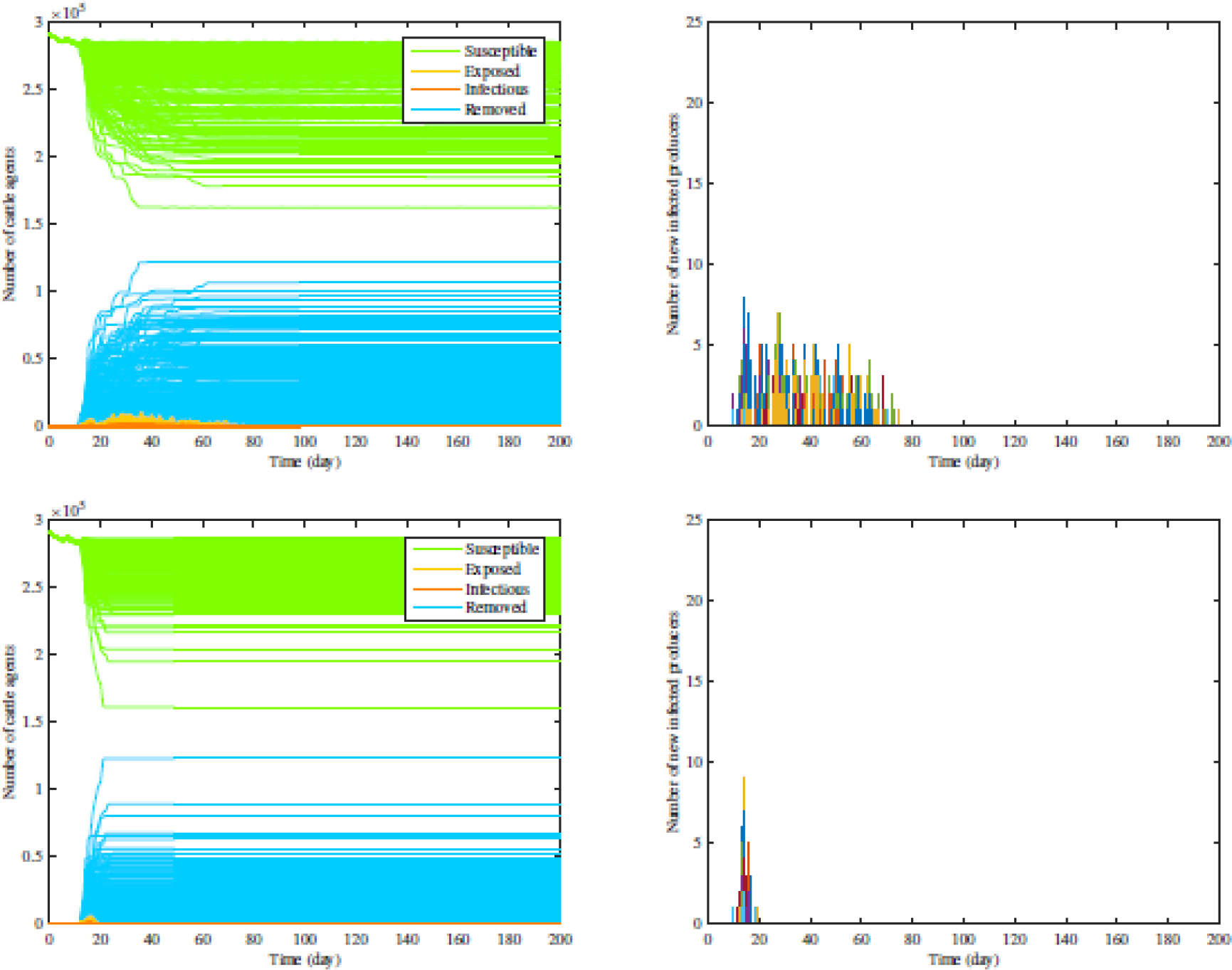
Comparison between scenarios 8 (top) and 9 (bottom).

In the cattle supply chain, cattle flows are essentially driven by the normal operation of packers to meet steady demand, so the total number of cattle in the system during the simulation period is strongly affected by the packers’ operation time. With infected cattle coming in, the packer will go to the under control state after a contamination period, and then will stop receiving cattle from feedlots both inside and outside the region. As a result, feedlots will stop requesting cattle from other premises, which will impact the number of cattle stocker operations will request. Therefore, the system’s cattle flow quickly stops once packers are under control, affecting the total number of cattle agents in the system. As the producers and packers gain revenues by marketing their cattle or cattle-related products, the total number of cattle agents in the system, which include the cattle that are marketed and being prepared to be marketed, can be seen as a measure for the economic impact of the control strategies on the cattle industry. The total packer operating time is the sum of the four packers’ operating time during the 200-day simulation period. Distributions of the total number of cattle agents that exist in the system and total packer operating time are shown in Fig. 6.

**Fig 6.**
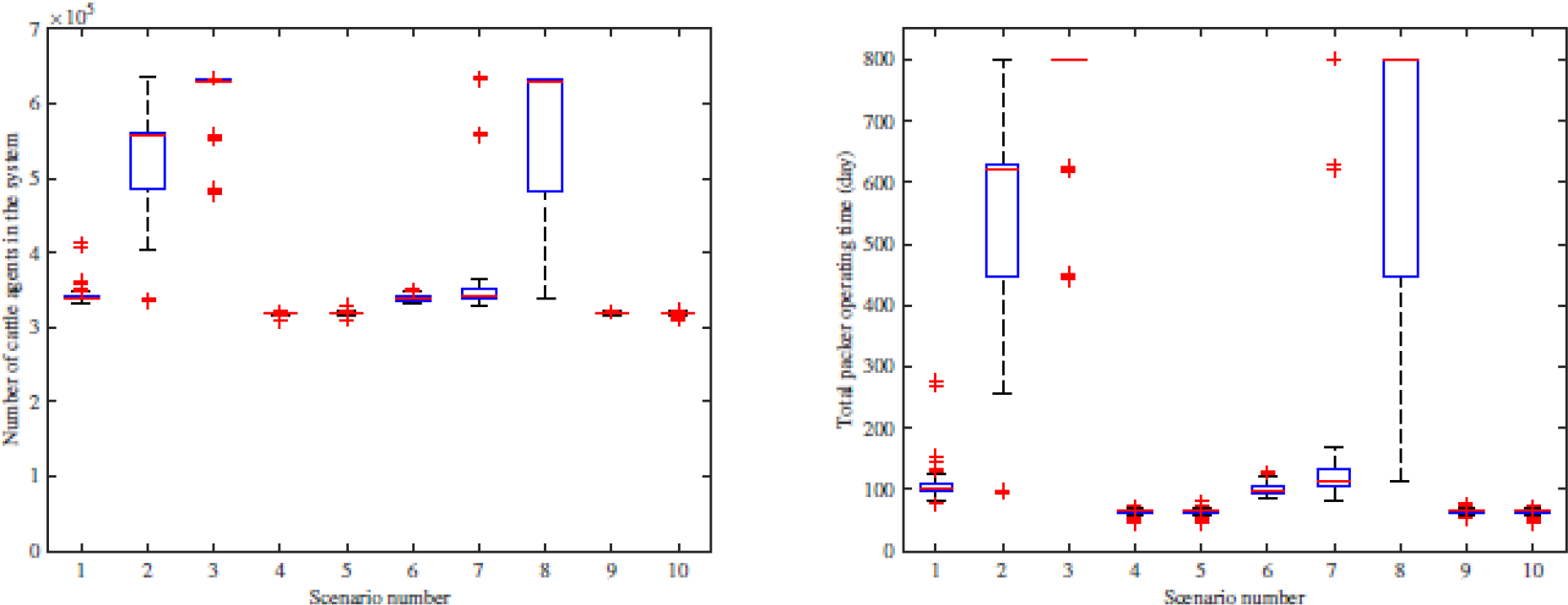
Distributions of the total number of cattle agents (left) and total packer operating time (right).

In Fig. 6, scenarios 3 and 8 result in the largest median number of cattle agents and packer operating time, followed by scenario 2 (producer isolation), and all the other scenarios are with a similar value. In scenarios 3 and 8 (information-sharing functionality enabled only on producers), the infected producers and their trading partner producers become isolated in the system quickly, but the packers can continue to request cattle from other uninfected producers, thereby sustaining regular operations for a longer time. Compared with scenario 1 (no farm-level control strategies implemented), the median number of cattle agents of scenario 2 is larger because producer isolation is already effective to contain the epidemic, resulting in less infected packers and an increased packer operation time. However, the infected producers will be under control more quickly in scenario 3 with information-sharing among producers, so packers can sustain regular operations for a longer time and the median number of cattle agents in scenario 3 is higher compared to scenario 2.

When information sharing is enabled for both producers and packers (scenarios 4 and 9), once infected feedlots occur, the four packers are much likely to have traded with these producers and will quickly transition to the under-control state, resulting in lower number of cattle agents over the system. Though information-sharing on all premises and region-wide movement ban (scenarios 4–5, 9–10) are both very effective regarding the epidemic containment, these two relatively protective strategies have a greater economic impact with a higher loss of cattle. It indicates that determining the appropriate control strategy is a tradeoff between multiple factors, including the economic effect and effectiveness of the epidemic control. Based on these simulations, placing control measures on packers has a large impact on the business discontinuity to the system based on decreased packer operating time and number of cattle agents. These control decisions need to be based on additional criteria not considered in the current setting, where a packer comes under control only because one of its trading partners is under control.

### 3.2 Sensitivity analysis

#### 3.2.1 Number of days to trace back and forward during the information-sharing process

The distribution of the number of infected producers under scenarios D&IN_B_P (information infrastructure enabled on producers) and D&IN_B_PP (information infrastructure enabled on producers and packers) is shown in Fig 7. The modeled outcomes are sensitive to the variation in the number of days to trace back and forward. For instance, a decrease from 14 days, in the baseline scenario D&IN_B_P (scenario 8), to 1 day, results in an increase of 21.4 times in the median number of infected producers. A small number of days to trace back and forward is not enough to contain the epidemic, but a larger value can lead to more premises under control, resulting in a larger economic impact. This implies that the number of days to trace back and forward is an important parameter during the information-sharing process and should be paid much attention in practice. Besides, the variation regarding the epidemic size in scenario D&IN_B_PP (scenario 9) is smaller than scenario D&IN_B_P, indicating that it is necessary to include the critical component packers into the information-sharing infrastructure.

**Fig 7.**
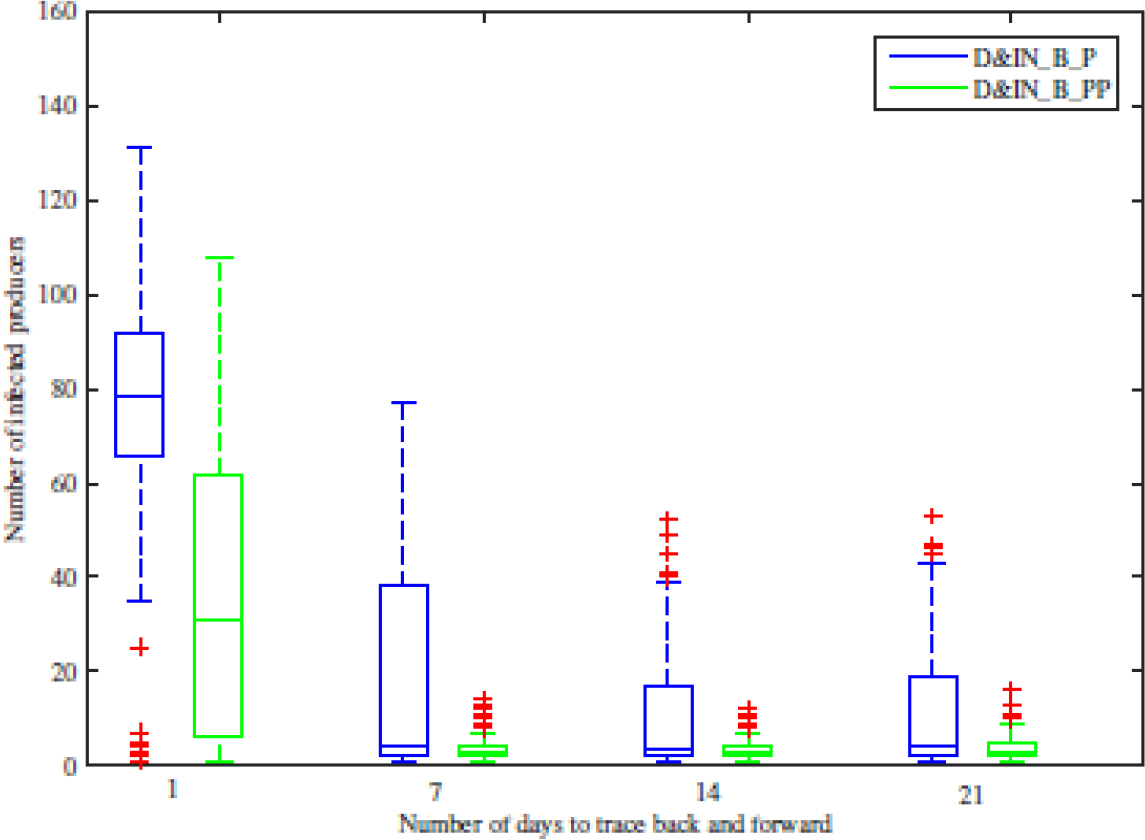
Sensitivity of the number of days to trace back and forward during information sharing.

#### 3.2.2 Indirect transmission probability and fomite survival time

The indirect transmission probability equals 0.15 in the baseline scenario, referring to the probability that a truck will become contaminated upon visiting an infected producer/packer, and the probability that contaminated truck will infect the subsequent producer/packer. Results in Fig. 8 are both sensitive to the indirect contact transmission probability and the fomite survival time for scenario D&IN_B (producer isolation). For example, an increase of indirect transmission probability from 0.15 to 0.2, the median number of infected producers changes from 82 to 112.

**Fig 8.**
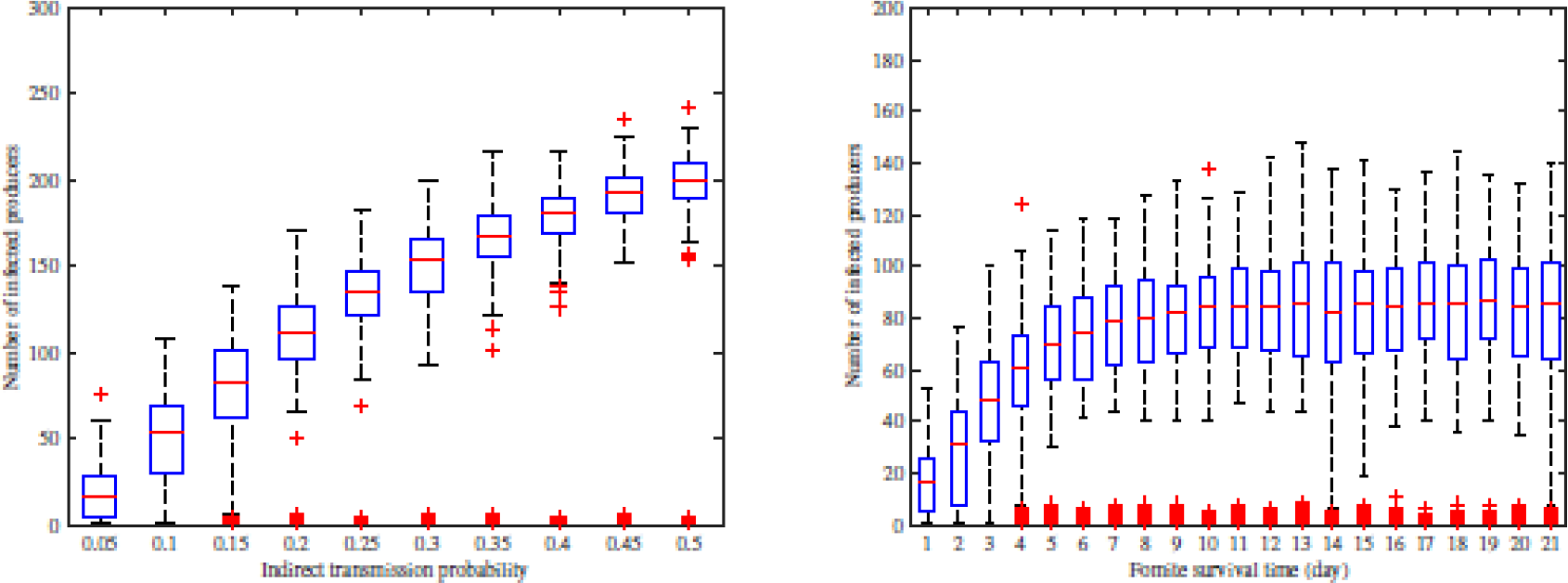
Sensitivity of indirect transmission probability and fomite survival time under scenario D&IN_B.

The sensitivity in outcomes to changes in indirect contact transmission probability and fomite survival time suggests that a more accurate estimation of these parameters related to indirect contact routes are critical to epidemic modeling. Since these two parameters are closely tied to the biosecurity status of cattle premises, these premises should be vigilant and follow guidelines for proper cleaning and disinfection of vehicles and cattle loading/receiving areas, such to limit the impact of indirect contacts on disease spread between premises.

#### 3.2.3 Impact of the cattle infectious period on epidemic sizes

Simulation results for variations in the parameter of the cattle infectious period associated with scenarios DI_B (direct contact only) and D&IN_B (direct and indirect contact) are shown in Fig. 9. Generally speaking, a longer cattle infectious period leads to a larger number of infected producers. For instance, variation in the cattle infectious period for scenario DI_B from 1 day to 9.2 days (Poisson distribution) results in changes in the median number of infected producers from 1 to 4.

**Fig 9.**
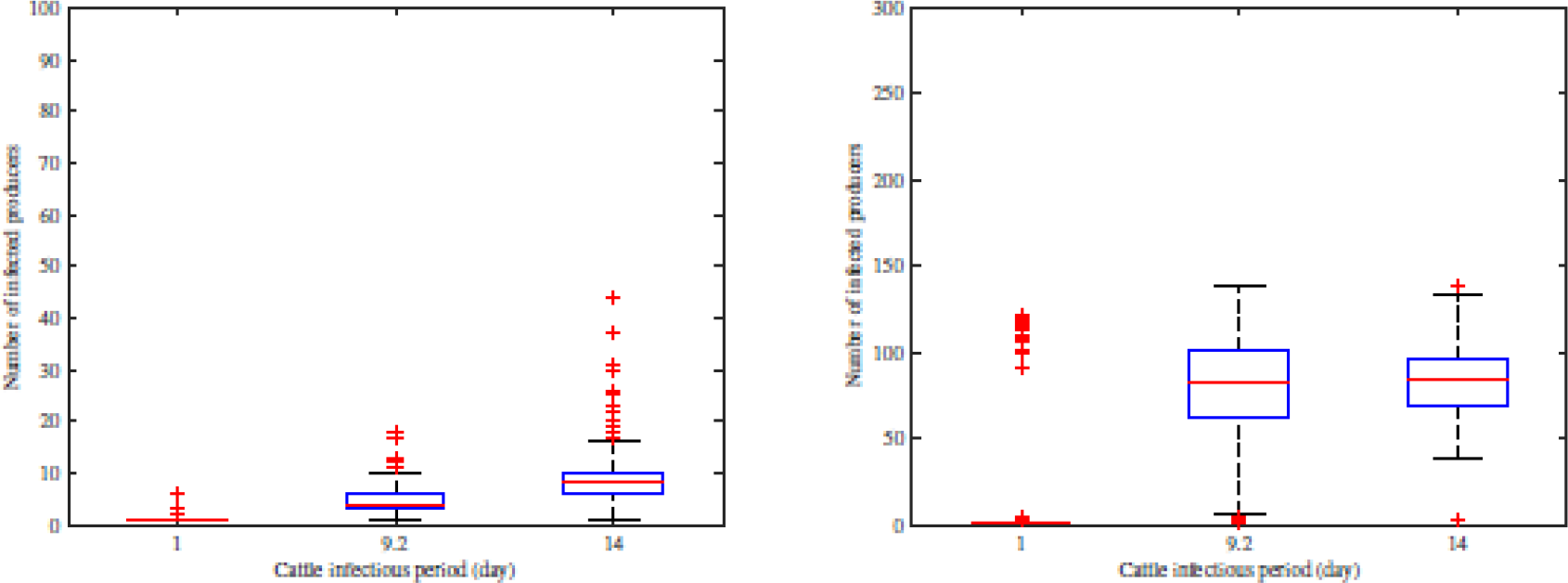
Sensitivity of the cattle infectious period under scenarios DI_B (left) and D&IN_B (right).

## 4. Conclusion

Based on the agent-based model developed in [38], which generates dynamic synthetic cattle and truck movement networks, this work analyzes the impact of truck contamination and information sharing on FMD transmission among premises in SW KS. Simulation results reveal the significant role truck movement (indirect contact) plays in further exacerbating the disease spread in the system, compared with equivalent scenarios that only consider animal movement (direct contact). For example, median [inter-quartile range] number of infected producers is 4 [3–6] vs. 82 [63–101.5] in baseline scenarios DI_B (direct contact only) and D&IN_B (direct and indirect contact) respectively. This highlights the need for a deeper understanding of indirect transmission mediated by fomites, and improved analyses or experiments need to be conducted to more accurately quantify fomite survival time and indirect contact transmission probability. Including information-sharing functionality on producers and packers can dramatically improve the system resilience against epidemics; for instance, the median number of infected producers in scenario D&IN_B (producer isolation) reduces by 96.3% in scenario D&IN_B_PP (information infrastructure enabled on producers and packers). Policymakers may focus more on promoting the development of new mitigation strategies informed by information sharing based on novel cyber-infrastructure for rapid detection and containment. Particularly, the number of days to trace back and forward during information sharing has a significant influence on the simulation outcome and is worth further studying considering the economic impacts of the related mitigation strategies.

Finally, it should be noted that we include only truck movement and ignore other forms of indirect contact, such as the movement of personnel or equipment. In the current model, producers/packers randomly select other producers to trade cattle, and future work can analyze the disease spread under various contact network structures by altering the current trade pattern. For simplicity, we assume that once packers are under control state, they stop requesting cattle and do not go back to normal operations during the epidemic period, while this may not be what happens in practice. Future work may impart greater realism to the packer agent and further analyze the economic impact caused by different control strategies.

## Acknowledgements

None

